# Lipid droplet biogenesis is driven by liquid-liquid phase separation

**DOI:** 10.1101/777466

**Authors:** Valeria Zoni, Rasha Khaddaj, Pablo Campomanes, Abdou Rachid Thiam, Roger Schneiter, Stefano Vanni

**Affiliations:** University of Fribourg, Department of Biology, 1700 Fribourg, Switzerland; Laboratoire de Physique de l’École Normale Supérieure, ENS, Université PSL, CNRS, Sorbonne Université, Université de Paris, Paris, France

## Abstract

Cells store energy in the form of neutral lipids packaged into micrometer-sized organelles named lipid droplets (LD). These structures emerge from the endoplasmic reticulum (ER), but their biogenesis remains poorly understood. Using molecular simulations, we found that fat accumulation and LD formation are described by a liquid-liquid phase separation (LLPS) process. Within this framework, we could identify how ER membrane properties modulate LD formation, and we could directly test our computational predictions by combining yeast genetics with fluorescence microscopy. Our data suggest that the specific lipid composition of the ER together with its peculiar physical properties, such as low membrane tension and membrane curvature, promote the packaging of neutral lipids into LD, preventing their accumulation in the ER membrane. Our results provide a new conceptual understanding of LD biogenesis in the context of ER homeostasis and function.

## Introduction

Lipid droplets (LD) are ubiquitous intracellular organelles that consist of a core of neutral lipids (NL), mostly triglycerides (TG) and sterol esters, surrounded by a phospholipid monolayer(*1*). Because of this unique composition, they are the cellular sites responsible for energy and lipid storage and they play a central role in lipid and cellular metabolism(*1–4*).

LD emerge from the endoplasmic reticulum (ER), where the NL constituting them are synthetized by acyltransferases that are essential for LD formation(*5*). These early LD have been observed with electron microscopy (EM), showing an oblate lens-like structure with diameters of 40-60 nm(*6*), and recent experiments indicate that they form at ER sites marked by the protein seipin(*7*), in combination with promethin (LDAF1)(*8*). However, because of important challenges in reconstituting *in vitro* the machinery of LD formation, the early molecular steps of LD biogenesis, remain largely unexplored. The mechanistic understanding of these steps(*3, 4, 9–14*) is currently based on the extrapolation of sparse experimental observations in reconstituted systems, where micrometric oil blisters have been observed, during the formation of black lipid membranes(*15, 16*) or in droplet-embedded vesicle systems (DEV)(*17*), suggesting that LD can spontaneously assemble following the accumulation of NL in between the bilayer leaflets. However, all these experiments lack the ability to continuously modulate and monitor the NL-to-phospholipids (PL) ratio during the process of LD formation, *i.e.* for increasing values of the NL-to-PL ratio as in cellular conditions, and it is thus difficult to establish the key factors controlling LD assembly based exclusively on information from reconstituted systems.

Molecular dynamics (MD) simulations represent a good alternative strategy to investigate the early stages of LD formation. This approach has been used to investigate the formation of oil blisters at two different NL-to-PL ratios, and it was found that NL blister formation takes place when the NL-to-PL ratio exceeds a certain threshold(*18*). Taken together, all these observations have led to a model that posits that LD form from the spontaneous accumulation of NL between the two leaflets of the ER into nanometer-sized lipid lenses at sites marked by the protein seipin, before a subsequent maturation that involves LD growth and budding(*3, 4, 9–14*).

Deletion of seipin, however, does not abolish LD formation, and, for what concerns the early stages of LD biogenesis, rather results in delayed LD formation and accumulation of NL in the ER(*19*). At the same time, deletion of other proteins with very different functions, such as the lipid phosphatase Pah1(*20*) or the membrane-shaping protein Pex30 together with seipin(*21*), results in a more pronounced phenotype, with the majority of cells completely lacking LD and with NL strongly accumulating in the ER(*20*) or in non-native NL-enriched membraneous structures(*21*). These observations reveal that LD formation is highly sensitive to the membrane environment, and they indicate that while seipin functions as the main ER nucleation seed for LD biogenesis, other mechanisms are likely to significantly contribute to the energetics of TG accumulation. From a conceptual point of view, the coexistence of NL in both the ER membrane as well as in nucleated LD that is observed in different non-physiological conditions(*19–21*), suggests that the process could be described in the framework of a liquid-liquid phase separation process (LLPS), a framework that has recently emerged as a key self-organizing principle for intracellular compartmentalization for membrane-less organelles in the cytosol(*22*). In the context of LD, nucleated fat lenses would constitute the condensed phase, while “free” NL in the ER would constitute the diluted phase, with the ER membrane as the solvent.

Given the small length scales of the process and the difficulty of controlling the ratio between NL and PL in both cellular and reconstituted approaches, we resorted to *in silico* MD simulations to investigate how membrane parameters can modulate the mechanisms of LD biogenesis. We found that this process is a *bona fide* LLPS and we identified how specific membrane properties are crucial to promote the packaging of NL into oil blisters. In order to validate *in vivo* our computational predictions, we took advantage of yeast genetics and we studied LD formation in different mutants, where we could rationally modulate LD formation in agreement with our computational predictions. Our results pave the way for a new conceptual understanding of LD biogenesis in the context of ER homeostasis and function.

## Results

### Triglycerides display liquid-like behavior in oil blisters and in lipid bilayers

To investigate the mechanism of LD biogenesis *in silico*, we first opted to reconstitute and further clarify the nucleation process observed by Khandelia et al(*18*). For this, we built lipid bilayers with increasing concentrations of TG (from 2% to 10%), with TG molecules initially distributed randomly between the two leaflets of the membrane (Figure 1A). We then let the simulations run towards equilibrium, and we observed that lens formation took place starting at TG concentrations above 4%, in agreement with what was previously reported(*18*). In addition, we observed that lens formation is concentration-dependent, as in classical nucleation processes(*14*) (Figure 1B).

**Figure 1.**
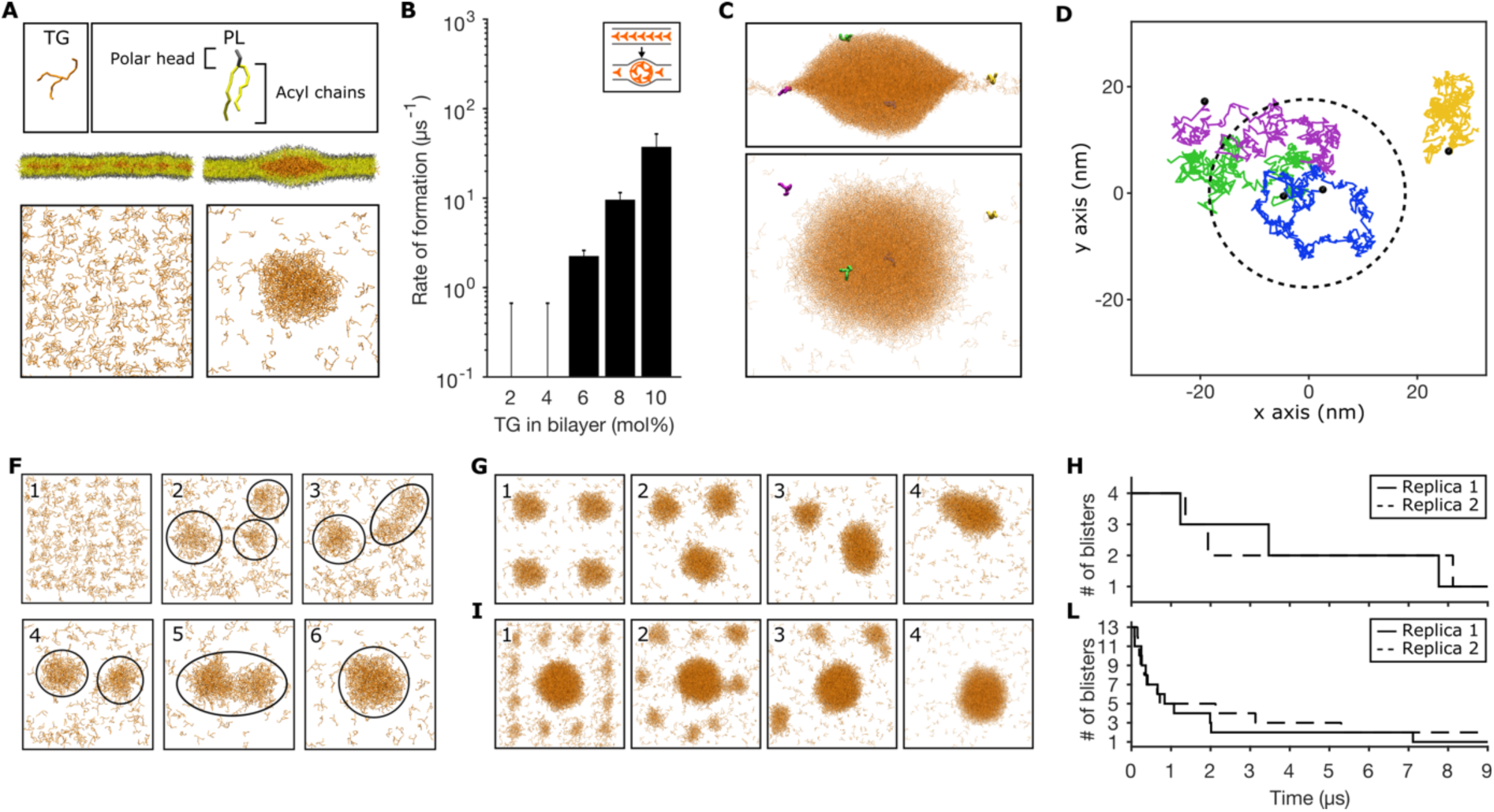
Triglycerides display liquid-like behavior in lipid bilayers and in oil blisters. (A) Setup used to investigate TG nucleation. (B) Rate of formation of oil lenses at different TG concentrations. (C) Side and top view of the starting configuration and (D) 2D single trajectories of 4 representative TG molecules: one that remains inside the TG lens (blue), one that remains free in the bilayer (yellow), one that exits (green) and one that enters (purple) the TG lens. (F,G,I) Coalescence of oil lenses in different MD simulations. Progressive numbers indicate the time evolution of the systems. (H,L) Number of lenses over time in the second (G) and third (I) coalescence *in silico* experiments.

Since both TG and lipid bilayers can adopt a liquid state at physiological temperature, we first tested whether TG molecules display liquid-like behavior in both the diluted and the condensed phase under the confinement imposed by the lipid bilayer. First, we computed TG diffusion in the bilayer, and we observed that TG freely diffuse in the lipid bilayer with a mean square displacement comparable to that of PL (Figure S1 and Table S1). Next, we observed that TG molecules are able to freely diffuse inside the blister, and can exchange between the two phases (Figure 1C,D, Movie S1).

**Table 1.**
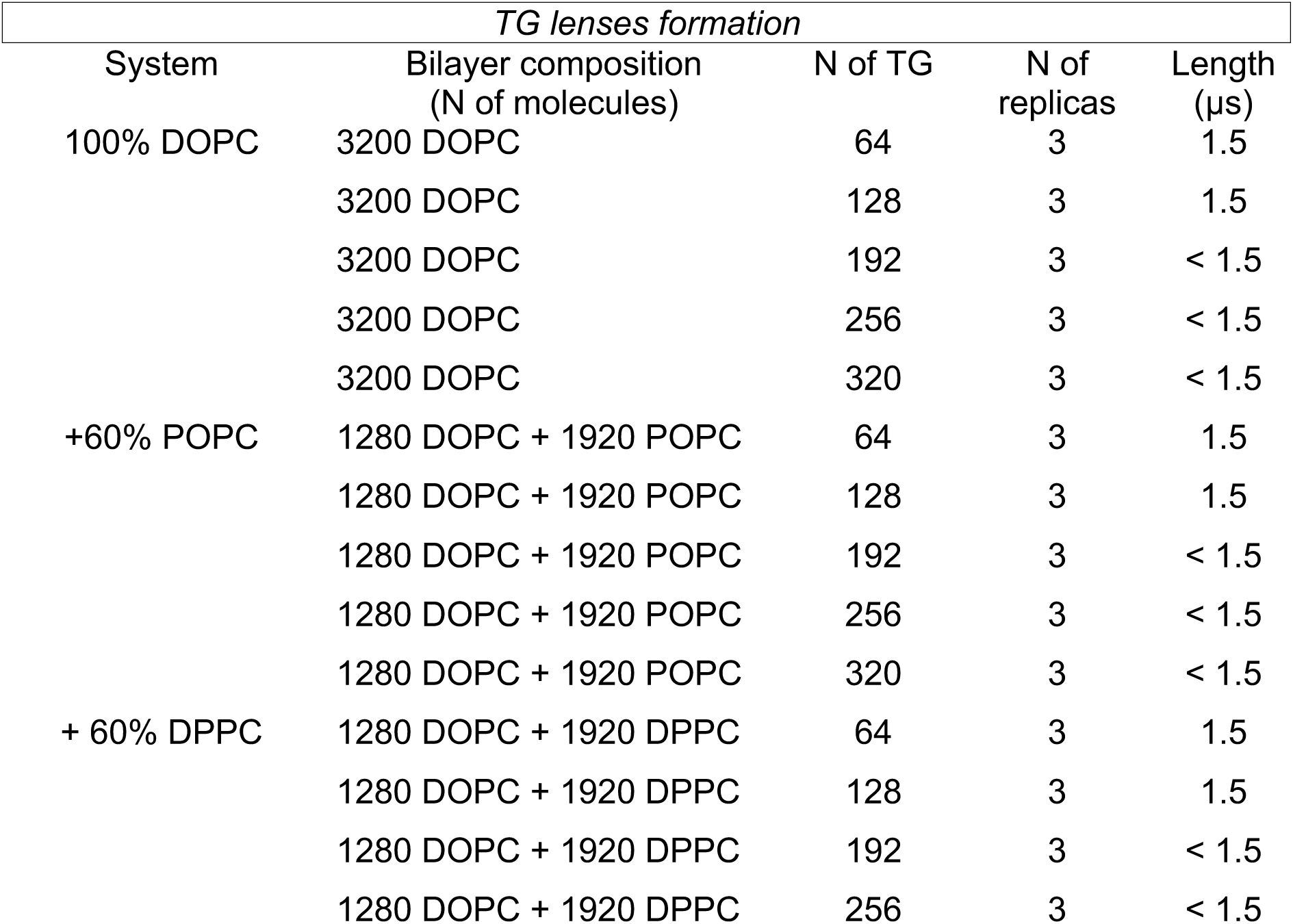
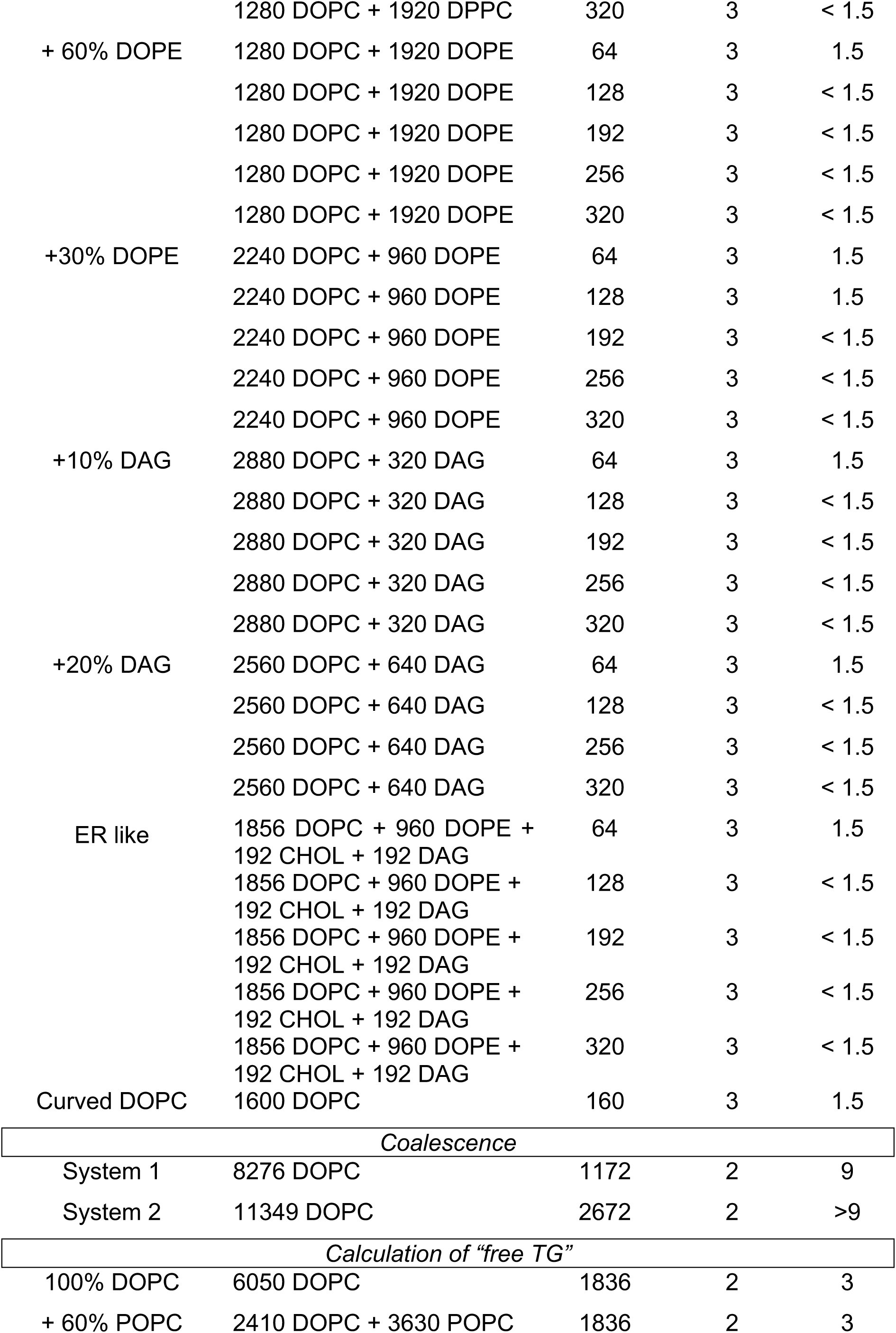
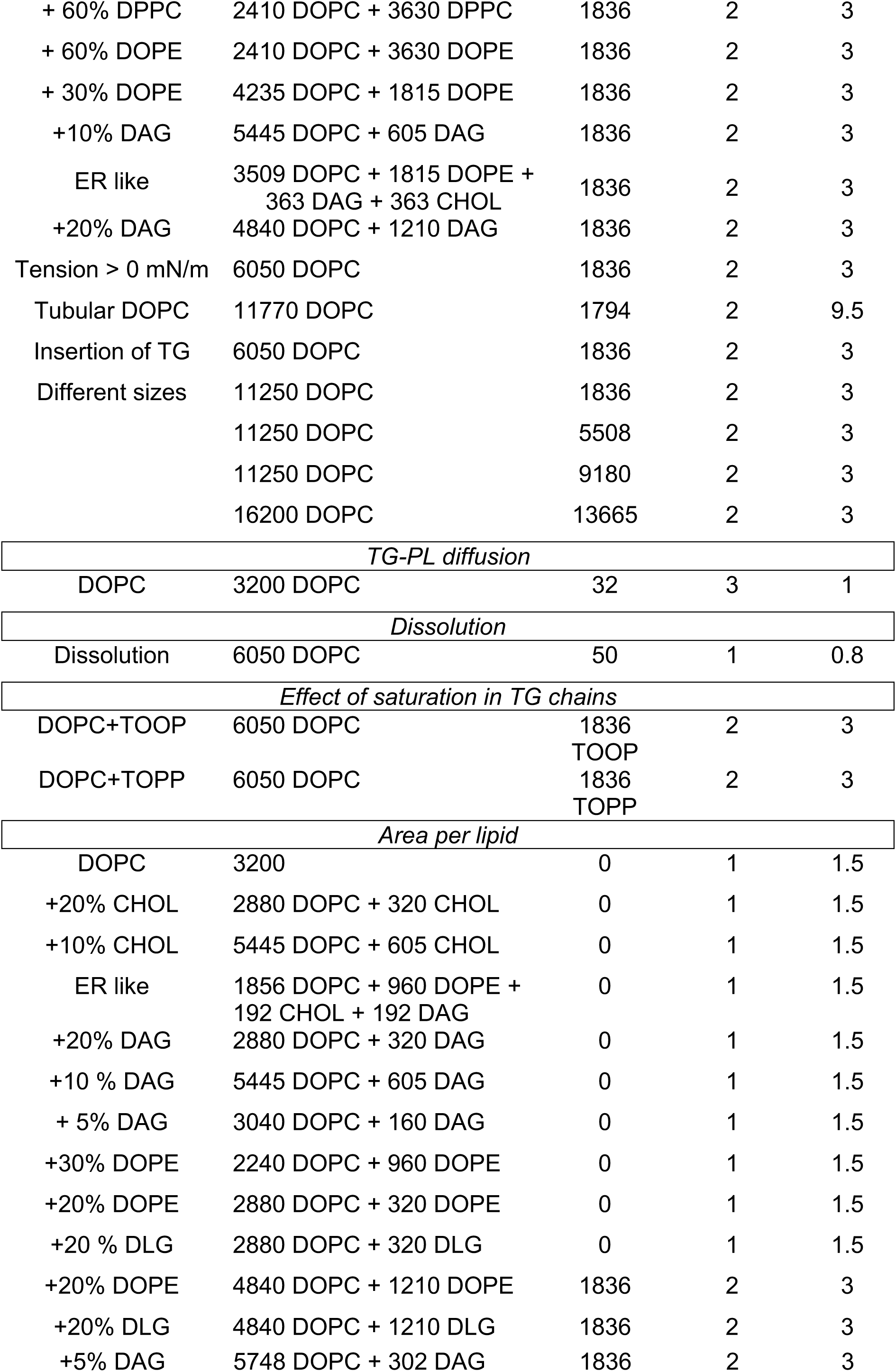
List of all the MD setups, with bilayer composition, number of TG molecules, number of replicas, and length of simulations.

Next, we tested whether oil blisters in bilayers can coalesce and fuse between them. To do so, we used three different computational assays: first, we observed that during spontaneous blister formation at high TG concentration (10%), multiple small lenses that initially formed in the simulation coalesced over time to a single one (Figure 1F). Second, we prepared a configuration with 4 identical lenses and observed their fusion to a single lens over time (Figure 1G,H). Third, we prepared a system with a big lens surrounded by 12 small lenses. Again, we observed their spontaneous coalescence, in this case accompanied by Ostwald ripening. (Figure 1I,L, Movie S2). Taken together, these results indicate that in our *in silico* reconstituted system, under the confinement imposed by the lipid bilayer, TG molecules display liquid-like behavior in both diluted and condensed phases. Our observation that TG molecules are liquid in the condensed phase is in agreement with the experimental observation that TG blisters in GUVs can spontaneously fuse and ripen(*7*).

### Oil-in-bilayer blister formation is a bona fide LLPS

One important aspect of LLPS is that the process must be driven by the equilibrium concentration (e.g. the chemical potential) of the solute in the diluted phase(*22*). For LD biogenesis, this would imply that while nucleation factors such as seipin could define the sites of LD formation, the thermodynamics of the process would be determined by the concentration of TG in the ER.

To test whether TG blister formation is a *bona fide* LLPS, we next investigated whether the mechanism is driven by the chemical potential of TG in the diluted phase. To do so, we first prepared a system that allowed us to estimate the concentration of TG molecules in a lipid bilayer by packing all TG molecules in a pre-formed lens inside a lipid bilayer, and we followed the time evolution of the concentration of TG molecules spreading into the bilayer (Figure 2A). After an initial equilibration, the system reached an equilibrium at the concentration of 1.1±0.1 mol% (Figure 2B), in close agreement with experimental values reported using capacitance measurements(*16*) or NMR(*23*).

**Figure 2.**
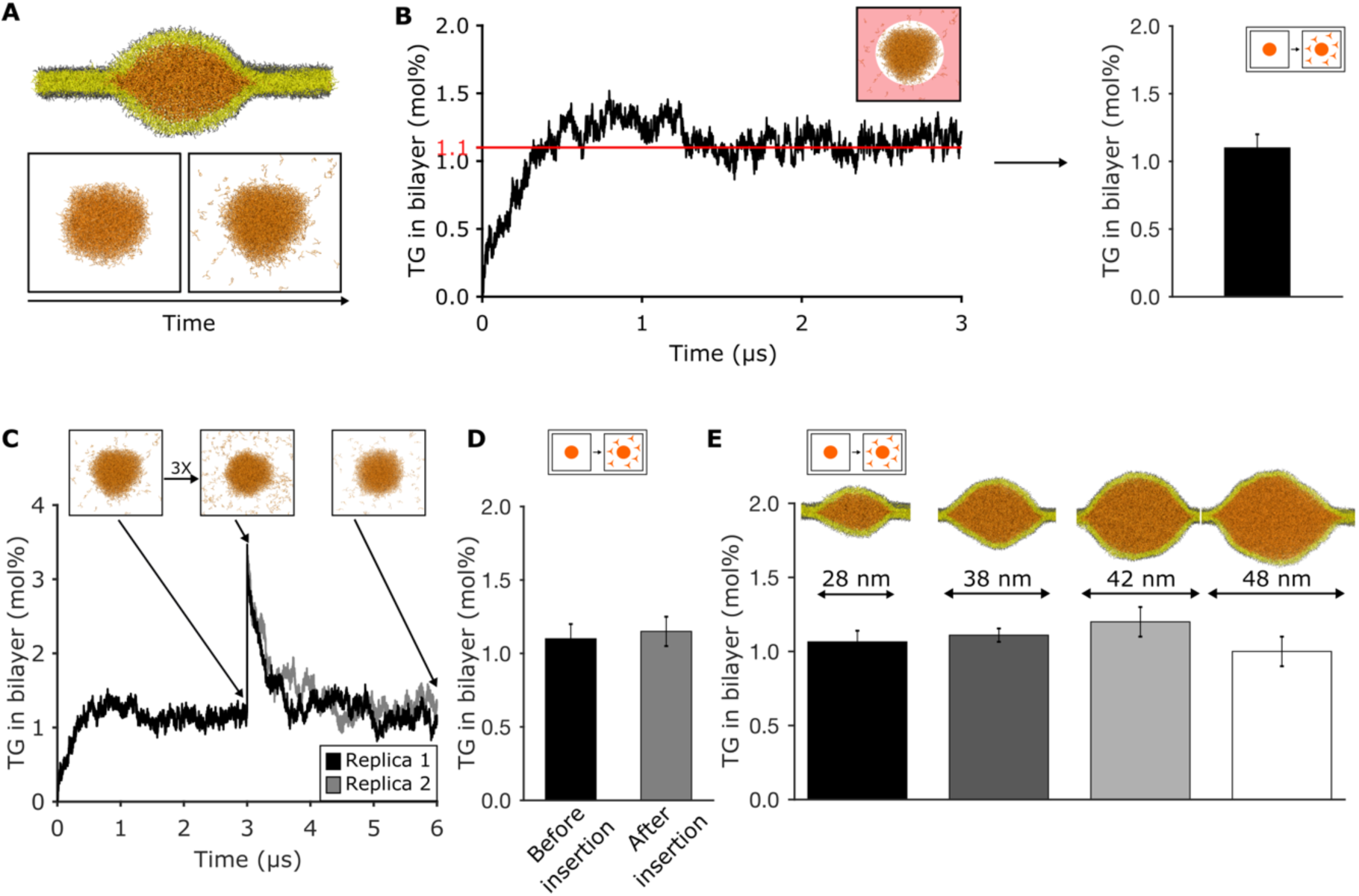
Oil blisters in lipid bilayers form via a liquid/liquid phase separation process. (A) Setup used to measure the amount of diluted TG via spontaneous diffusion from a pre-formed blister and (B) quantification over time. The area in which TG is assumed as diluted is highlighted in light red. (C) Time evolution of the percentage of diluted TG inside the bilayer. The injection of additional TG molecules was performed after 3 μs of dynamics. (D) Comparison between the values of diluted TG before and after the insertion of TG in the bilayer. (E) Quantification of diluted TG in oil blisters of different sizes.

To test whether the TG concentration in the bilayer is the driving force of the process, we used three different approaches. First, we arbitrarily increased the concentration of the diluted TG in the bilayer by computationally “injecting” new TG molecules in an equilibrated system (Figure 2C). When the system reached equilibrium, all excess TG molecules translocated to the oil lens, with the bilayer TG concentration returning to its initial value after few microseconds (Figure 2C,D).

Second, we prepared oil lenses of different sizes compatible with the ones observed using EM(*6*), ranging from 25 to 50 nm in diameter (Figure 2E). We observed that the concentration of equilibrium TG in the bilayer is independent of the blister size. Third, when TG blisters with a total TG concentration below the computed chemical potential (1.1% TG/PL) were simulated, this led to blister dissolution (Movie S3). Taken together, our results indicate that indeed oil blister formation in lipid bilayers can be considered as a *bona fide* LLPS.

### Membrane physical properties modulate oil-in-bilayer LLPS

LLPS is a ubiquitous cellular strategy to achieve membrane-free intracellular compartmentalization(*22*). The underlying molecular mechanisms governing the precise regulation of these processes in cellular systems, however, remain mostly elusive(*22*). Having established that oil blister formation *in silico* proceeds through a LLPS, we hypothesized that in the specific case of intracellular LD, solvent properties (i.e. characteristics of the ER membrane) could play a major role in modulating the thermodynamics of LD formation.

We therefore tested *in silico* whether membrane properties could modulate the propensity for blister formation and, concomitantly, the concentration of TG molecules residing in the lipid bilayer. We first investigated whether physical properties, such as membrane tension and membrane curvature, might modulate this mechanism. Of note, the ER, unlike other organelles, is characterized by low membrane tension(*24*) and extensive membrane curvature(*25*), that is apparent in its characteristic tubular and sheets structure.

By modeling LD formation at different surface tensions, we observed that increasing membrane tension (Figure 3A) increases the amount of diluted TG in bilayers (Figure 3B), and thus antagonizes blister formation, in agreement with *in vitro* experiments(*17*). Membrane curvature (Figure 3C), on the other hand, did not appear to affect the chemical potential of TG molecules in the bilayer (Figure 3D). This observation appears to be at odds with the recent experimental observation of a prominent role of the membrane-shaping protein Pex30 in LD formation(*21, 26*). However, in agreement with our data, when the major ER-shaping proteins, such as the reticulons proteins Rtn1 and Yop1, were deleted in yeast, this did not result in the accumulation of NL in the ER membrane(*21*). This suggest that the specific membrane remodeling effect of Pex30 is likely related to LD nucleation, as Pex30 marks the sites of LD formation together with seipin(*21*).

**Figure 3:**
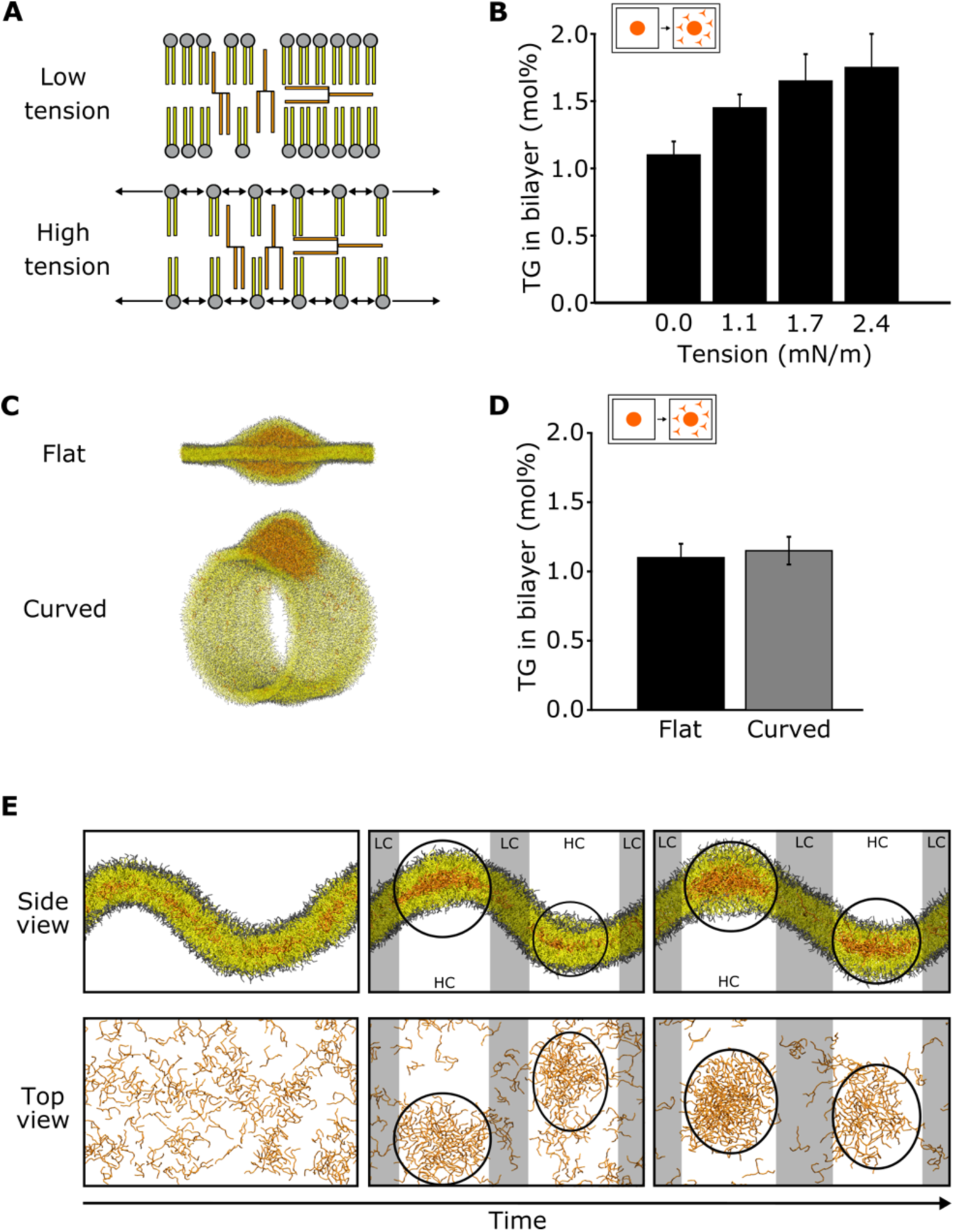
Membrane physical properties modulate LD biogenesis. (A) Cartoon of the TG-enriched bilayer at different tensions. (B) Percentage of diluted TG at different surface tensions. (C) MD snapshots of flat and curved systems used to quantify the percentage of diluted TG in flat and curved systems. (D) Percentage of diluted TG in flat and curved systems. (E) Representative snapshots of the simulations leading to blister formation in a curved system. Oil blisters are highlighted with black circles. Regions of high curvature (HC) are highlighted in white, while regions of low curvature (LC) are highlighted in gray.

To test this hypothesis, *i.e*. whether curvature affects the spatial localization rather than the energetics of droplet formation, we visually investigated the MD simulations of oil nucleation in lipid bilayers with coexisting flat and curved patches (Figure 3E). We invariantly observed that at high TG concentrations, when spontaneous formation of oil blisters took place, lens invariantly nucleated at sites of high curvature, remaining there throughout the entire simulation (Figure 3E).

### ER lipids modulate oil-in-bilayer LLPS

Next, we investigated the effect of PL on the process, with a specific focus on those that are relatively abundant in the ER, and namely those carrying unsaturated chains and ethanolamine as their headgroups (PE) (Figure 4)(*27, 28*). By preparing lipid bilayers with PL with increasing levels of saturated acyl-chains (Figure 4A), we observed that lipid unsaturation slightly promotes blister formation (Figure 4B) and, correspondingly, segregation of TG from the membrane (Figure 4C). Similarly, the presence of conical-shaped PE lipids (Figure 4D) facilitated nucleation of TG (Figure 4E) and decreased the equilibrium concentration of TG in the bilayer (Figure 4F).

**Figure 4.**
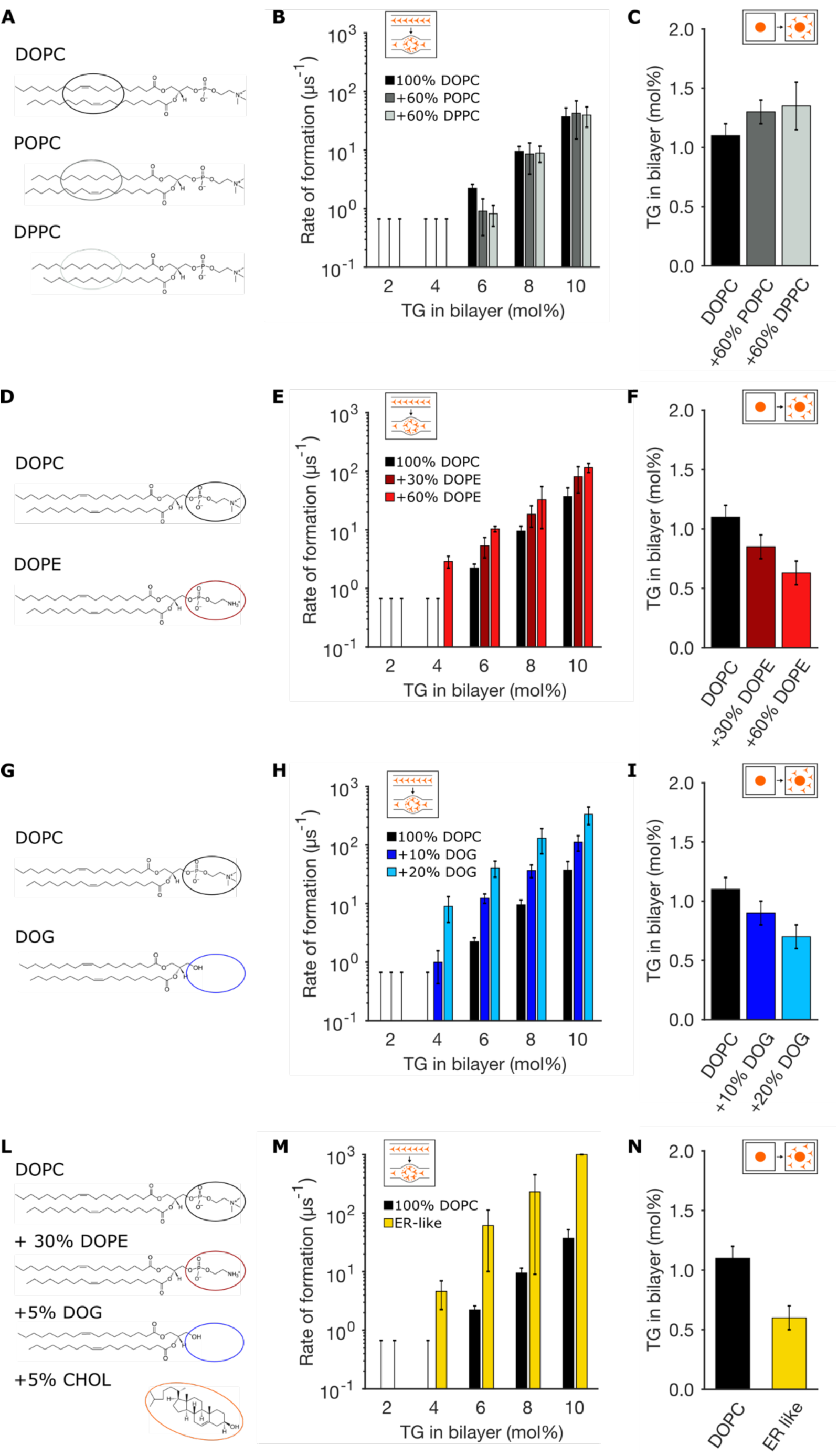
ER composition modulates TG aggregation. (A, D, G, L) Lipid compositions tested and chemical structures of the various lipids involved in the mixtures. (B, E, H, M) Corresponding rates of blister formation, and (C, F, I, N) percentage of diluted TG.

Next, we reasoned that at sites of LD formation, the lipid precursor of TG, diacylglycerol (DAG), could be present at higher concentration, and we sought to investigate the effect of increasing concentrations of DAG on LLPS (Figure 4G-I). Remarkably, we observed a pronounced effect towards blister formation, with even mild concentrations of DAG promoting phase separation at low (<4%) TG concentrations (Figure 4H).

Lastly, we prepared bilayers with a lipid mixture that approximates that of the ER (Figure 4L), by including PL unsaturation, high PE content, and low but non-negligible quantities of DAG and cholesterol(*28*). As it can be seen in Figure 4M,N, the effect of the various lipids adds up, resulting in a dramatic acceleration of the LLPS. Taken together, our results suggest that the ER membrane appears engineered to promote oil blister formation, allowing only very low concentration of diluted TG in the ER, possibly to reduce TG-induced lipotoxicity.

### Lipid remodeling alters the equilibrium between LD formation and accumulation of NL in the ER consistently with a LLPS framework

Our model suggest that LD formation is a LLPS that can be modulated by specific membrane properties. In particular, we observed *in silico* that membrane composition plays a major role, with unsaturated lipids, PE and DAG all contributing to promote oil blister formation. To test our theory *in vivo*, we resorted to fluorescence microscopy experiments in yeast, a model system that has been widely used to investigate LD biogenesis. We reasoned that if we could modulate the balance of NL between LD and the ER membrane (*i.e.* between the condensed and diluted phases in our LLPS framework) with independent approaches consistently with our *in silico* predictions, this would provide strong evidence in support of our theory.

As a first test, we focused on the role of the precursor of TG, DAG, as our simulations indicate a major effect of DAG in promoting oil blister formation (Figure 4G-I). In yeast, deletion of Pah1, the lipid phosphatase converting phosphatidic acid (PA) into DAG upstream of TG synthesis, interestingly results in a phenotype where NL are present in the cell at significant concentrations but fail to cluster into LD and rather accumulate in the ER(*20*), consistently with our *in silico* data.

To further test the effect of DAG concentration on LD biogenesis, we expressed a previously-characterized DAG sensor in the ER of yeast cells(*29*) This sensor is comprised of tandem C1 domains from human protein kinase D (C1a/b-PKD) fused to GFP, which in turn is fused to the transmembrane region of Ubc6, a tail anchored domain protein(*29*). With this probe, we could compare the localization of DAG in the ER of wild-type cells with that of cells in which NL fail to be packaged into LD and are rather spread over the entire ER, such as those where TG is the only NL and Pah1 is deleted (Figure 5A,B). In this condition, while both DAG and TG are present in the ER(*20*), DAG adopts a slightly more punctate distribution in wild type cells in comparison with *pah1Δ* mutant cells (Figure 5A,B). The localization of TG, as shown by BODIPY fluorescence, is strikingly different in the two cases (Figure 5A,B), with a punctate localization in the wild type, indicating the presence of LD, and a diffuse ER staining in the case of *pah1Δ* mutant cells. These data suggest that increased DAG local concentration at specific ER sites might promote LD formation, in agreement with our *in silico* simulations.

**Figure 5:**
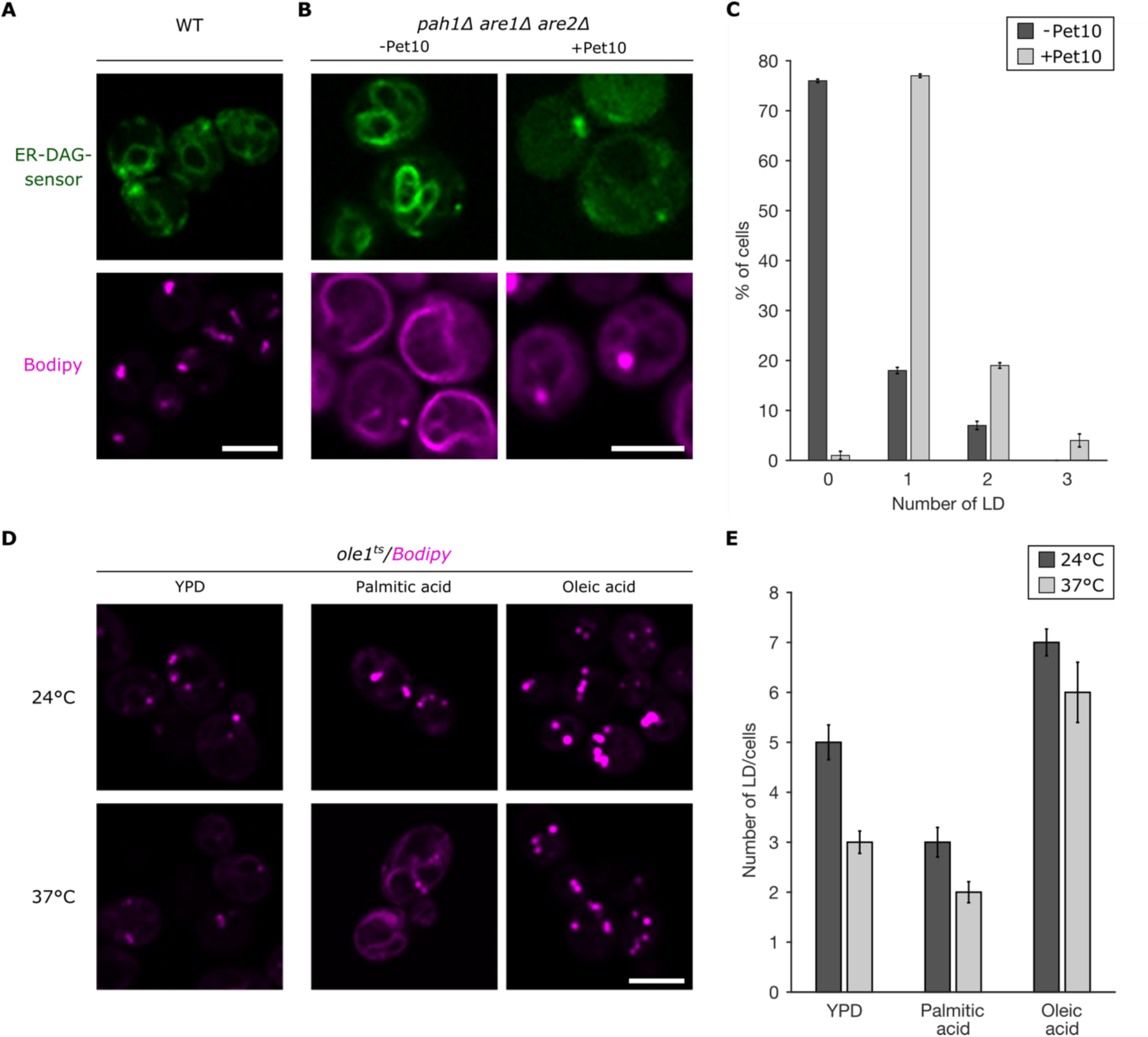
ER membrane remodeling modulates LD formation. (A) Fluorescence microscopy images of WT and (B) *pah1Δare1Δare2Δ* (with or without overexpression of Pet10) yeast cells. Top panels: ER-DAG sensor staining. Bottom panels: NL are stained with BODIPY. (C) Quantification of the number of LD per cell (*n*=100) with and without Pet10 overexpression. (D) Fluorescence microscopy images of temperature-sensitive *Ole1* at 24°C (top) and 37°C (bottom). Cells are stained with the NL marker BODIPY. Strains were cultivated in YP-rich medium (Left column) or upon supplementation of palmitic acid (middle) or oleic acid (right). (E) Quantification of the number of LD per cell (n=20) in the different conditions shown in (D): YPD, palmitic acid, oleic acid.

To further test this hypothesis, we investigated whether DAG accumulation could restore LD formation in the *pah1Δ* mutant. To do so, we monitored DAG localization upon overexpression of Pet10, the yeast perilipin(*30*), since perilipins have been suggested to bind DAG in the ER(*31*). We indeed observed that, upon overexpression of Pet10, DAG adopts a non-uniform localization in the ER and that, correspondingly, TG aggregate in LD (Figure 5B). By counting the number of LD per cell in the absence or presence of Pet10 overexpression, we quantified that while more than 70% of cells are devoid of LD in the absence of Pet10 overexpression, almost all cells have at least one LD after DAG clustering induced by Pet10 (Figure 5C). These observations suggest that, in a cellular context, an elevated local concentration of DAG, possibly promoted by Pah1 activity at specific ER sites(*32*), can drive the packaging of TG into LD, promoting their biogenesis, consistently with our simulation data.

Next, we investigated the equilibrium between TG in the ER *vs* LD in yeast cells with an altered ratio between saturated and unsaturated fatty acids, as our simulations suggested (Figure 4A-C) that saturated PL decrease LD formation and promote accumulation of TG in the lipid bilayer. For this we used a temperature-sensitive allele of *Ole1*, a gene that is essential for production of monounsaturated fatty acids as it expresses the lone delta-9 desaturase enzyme in yeast(*33*). This temperature-sensitive allele results in intact proteins at permissive temperature (24°C) but inactive proteins at non-permissive temperature (37°C) and, concomitantly, to a major decrease in the ratio between unsaturated and saturated fatty acids in the cell(*34*). As fatty acids are incorporated in both PL and TG, we also investigated *in silico* the effect of increasing acyl chain saturation in TG, and we found that, like for PL, increasing TG saturation antagonizes LD formation and results in TG accumulation in the bilayer (Figure S2).

Using fluorescence microscopy, we observed Bodipy-positive ER structures at the non-permissive temperature, indicating that the amount of NL present in the ER increases as a result of the increase in saturated cellular fatty acids (Figure 5D). This enrichment was further promoted by supplementing the cells with the saturated palmitic acid (Figure 5D), while it was entirely rescued by supplementing the cells with the unsaturated oleic acid (Figure 5D). To further quantify this behavior, we opted to count the number of LD per cell in the different conditions, and we could confirm that an increase in the ratio between saturated and unsaturated fatty acids leads to a decrease in the propensity of LD formation, in agreement with our *in silico* data (Figure 5E).

## Discussion

The current model of LD formation posits that after synthesis in the ER membrane, TG bind to the oligomeric seipin-promethin (LDAF1) complex, that then acts as a nucleation seed for the formation of a nascent TG lens. When seipin is depleted, LD can still form, but their size distribution is misregulated and LD formation is delayed(*19*), resulting in a mild accumulation of TG in the ER(*35*). Remarkably, this accumulation of TG in the ER is dramatically enhanced when, in yeast, the seipin homologue Fld1 is deleted in combination with the ER membrane shaping protein Pex30(*21*), or if the lipid phosphatase responsible for converting PA into DAG, Pah1, is genetically removed(*20*). On the one hand, these experiments suggest that a detectable accumulation of TG in the ER is a hallmark of non-physiological conditions, under which proper LD formation does not take place; on the other hand, they suggest that seipin-independent ER properties, such as DAG synthesis or ER morphology, can significantly alter the propensity of LD formation.

Our data indicate that these experimental observations can be interpreted in molecular terms by considering LD formation and TG accumulation as a liquid/liquid phase separation, with the condensed phase being the nucleated nascent LD and the diluted phase being the TG-enriched ER membrane. This framework, however, not only helps providing a molecular interpretation to the existing experiments, but also furthers our understanding of the energetics underlying the process of TG accumulation(*3, 4, 10–14, 36*).

In a LLPS, the driving force behind the process is the chemical potential of the solute in the diluted phase, in our case the concentration of TG in the ER membrane. In simple terms, this means that no LD formation can take place when the concentration of TG in the ER is below the critical concentration, while LD formation takes place spontaneously when the concentration of TG exceeds that threshold, albeit with a kinetics that depend on the height of the nucleation barrier.

This has important implications for both ER homeostasis and LD biogenesis and growth. First, this suggests that the amount of TG in the ER at equilibrium is not zero. This has potential implications in the catabolism and homeostasis of TG as it opens the possibility that lipases or lipid transport proteins might interact with TG directly from the ER, rather than upon binding to LD. Second, LLPS provides an efficient mechanism to promote and sustain LD growth, as once the nascent LD is formed, all TG in excess of its equilibrium chemical potential in the ER will flow to the existing LD without need for additional external energy, as long as the protein machinery around the LD-ER contact site does not prevent free diffusion of TG. This is particularly important as pre-existing LD that are resistant to starvation have been observed, for example in COS-1 cells (*32*).

Our data suggest that specific properties of the ER membrane, and namely low membrane tension and abundance in both mono-unsaturated lipids and phosphatidyl-ethanolamine(*24, 28*) contribute to keep TG in the ER at very low concentrations, thus not only promoting their packaging into LD, but also potentially preventing their lipotoxicity(*37*). On the other hand, we surprisingly found that membrane curvature, that is ubiquitous in the ER membrane(*25*), does not affect TG concentration in lipid bilayers. However, we identified that membrane curvature plays a role in LD nucleation, providing a mechanistic explanation of why only membrane-shaping proteins that colocalize with the marker of LD biogenesis seipin, such as Pex30, alter LD formation propensity, while membrane-shaping proteins that are ubiquitously found in ER tubules, such as Reticulons or Yop1, do not affect LD biogenesis(*21*). This observation also suggests that colocalization between TG synthesis and LD nucleation might provide an energetically efficient strategy to accelerate the process of LD formation.

By varying the lipid composition in the ER membrane of yeast by means of genetic strategies, we could alter LD formation consistently with our computational predictions. We speculate that similar variations in ER lipid composition could also take place in the cell in response to different stimuli. As a notable example, we found that the presence of the TG precursor, DAG, promotes LD formation. Thus, during PA to DAG conversion, not only TG synthesis increases, as more substrate becomes available, but also the physical prowess of the ER membrane for TG packaging increases. Similar feedback loops can arise in response to environmental variations, with cells potentially adjusting the local composition of the ER to, for example, promote or delay LD formation in response to the intake of different nutrients. Interestingly, we observed that an increase in the amount of saturated fatty acids challenges LD formation in yeast cells, rather promoting TG accumulation in the ER. This could potentially provide a new molecular explanation of the elevated toxicity of saturated fatty acids that is independent to the efficiency of their incorporation into TG(*37–39*).

A comprehensive experimental determination of the key ER factors that modulate LD biogenesis, however, remains extremely challenging, for at least two major reasons. First, it would require precise measurements of local lipid abundance and dynamics. Second, even in simple model systems such as those we have investigated *in silico*, we observed that basically every property of lipid bilayers (e.g. tension, curvature, lipid composition) can, in one way or another, affect LD formation. However, a few major trends emerge from the analysis of the MD simulations. First, lipid shape, or intrinsic curvature, seems to play a significant role, as lipids with conical shape such as unsaturated-PC, PE or DAG seem to promote LD formation. However, this property is not sufficient to describe the propensity for LD formation, as, for example, this observation does not explain the behavior observed when varying membrane surface tension. A better correlation can be established between the propensity of LD formation and the effective area per lipid in TG-free lipid bilayers (Figure S3), as this explains not only the observed behavior for conical-shaped lipids, but also the decreased propensity for LD formation that we observed when increasing membrane tension. This suggests a possible excluded-volume effect as the explanation for the behavior we observed: when lipids are tightly packed, it is energetically unfavorable for TG to reside in the interior of the bilayer and, as such, their separation into nascent oil lenses is favored. On the other hand, when the lipids are loosely packed, TG can occupy the interior of the bilayer at much higher concentrations, and possibly even help relieve the stress arising from such an inadequate spatial arrangement, as in the case of high values of membrane tension. As physiological membranes are extremely complex, however, it remains unclear whether the combination of multiple factors (e.g. acyl chain length, membrane tension, curvature, domains,….) has synergistic or cooperative effects toward LD formation and what are ultimately the main factors shaping this process.

Our results, however, provide a new conceptual view on the mechanism of LD biogenesis and fat accumulation, whereby not only local events taking place at the specific sites of LD formation, but also more global properties of the surrounding ER membrane play a major role in the overall regulation of this process. On the one hand, these properties are easier to measure, for example by means of genetic manipulation of lipid composition coupled with lipid imaging techniques, than local properties such as LD surface or line tension. On the other hand, our framework suggests that physiological variations in ER lipid composition at sites spatially distinct from LD, for example by means of lipid transport or establishment of contact sites with other organelles, might ultimately provide an alternative cellular route to modulate intracellular fat accumulation.

## Materials and Methods

### Molecular dynamics (MD) simulations

All MD simulations were performed using the software LAMMPS(*40*) and employing the Shinoda-Devane-Klein (SDK) force field(*41–44*). Parameters for TG were taken from (*42*) and triolein was chosen as the model TG, unless explicitly otherwise stated (Figure S2). Combination rules, as described in (*45*), were used to derive non-bonded interactions of DAG with DOPE and cholesterol. Initial configurations and input files were obtained through conversion of atomistic snapshots using the CG-it software (https://github.com/CG-it/).

In the simulations, temperature and pressure were controlled via a Nosé-Hoover thermostat(*46*) and barostat(*47–49*): target temperature was 310K and average pressure was 1 atm. The lateral *xy* dimensions were coupled, while the *z* dimension was allowed to fluctuate independently. Temperature was dumped every 0.4 ps, while pressure every 1 ps. Linear and angular momenta were removed every 100 timesteps. In simulations performed at non-zero surface tension, the *xy* dimensions were kept fixed with no pressure-coupling. Van der Waals and electrostatic interactions were truncated at 1.5 nm. Long-range electrostatics beyond this cutoff were computed using the particle-particle-particle-mesh (PPPM) solver, with an RMS force error of 10^-^⁵ kcal mol^-^¹ Å^-^¹ and order 3. In all CG-SDK systems, a time step of 20 fs was used, except for bilayers containing cholesterol, where a time step of 10 fs was used.

### MD systems setup

In order to study the formation of TG lenses (Figure 1A-B), bilayers containing different concentrations of TG were employed. To build these systems, boxes consisting of 50 phospholipid (PL) molecules and a variable number of TG molecules, to see the effect of different TG concentrations on lens formation, were initially prepared. The two PL monolayers were displaced along the z-axis in order to allow the insertion of the TG molecules and avoid steric clashes due to bad contacts or overlapping molecules. The TG molecules were subsequently randomly placed between the two monolayers. Boxes were then replicated 8 times along the x and y axes. Every system finally included a total of 3200 PL molecules. Different bilayer compositions were tested, as reported in Table 1. Simulations were run until spontaneous lens formation (see *MD simulations analysis* for details) or for a total of 1.5 μs per run if no spontaneous lens formation was observed (Table 1).

To investigate the coalescence of oil blisters, two additional systems were built. The first one (Figure 1G) was created from the last snapshot of the simulation with DOPC and 10% TG (radius 7 nm, Figure 1F), which was replicated 4 times along the x and y dimensions. To build the second, we merged the last frame of the simulation carried out to calculate the amount of “free TG” (see below) with that of the simulation performed to investigate TG lens formation (DOPC and 6% TG). This combination led to a big lens (14 nm radius) in the center surrounded by 12 smaller lenses, each of them with a radius of 5 nm, (Figure 1I).

The curved systems (Figure 3E) were created by building bilayers of 1600 DOPC molecules containing 160 TG molecules uniformly distributed. After equilibration, a force along the *x* direction was applied and then, during production, the *x* and *y* dimensions were kept fixed, as described in (*50*).

To calculate the concentration of diluted TG in the bilayer (Figure 2A-B and Figure 4C,F,I,N), systems were formed by positioning a lens of 1836 TG molecules between two monolayers with 3025 DOPC lipids each, as in (*17*). We also created bigger systems, with 12250 lipids and 1836, 5508, and 9180 TG molecules or with 16200 DOPC and 13665 TG, in order to study the size independence on the concentration of diluted TG in the bilayer at equilibrium (Figure 2E). All bilayer compositions described in Table 1 were tested.

To simulate systems at non-zero surface tension (Figure 3B), the system above (consisting of 1836 TG molecules between two monolayers with 3025 DOPC lipids each) was initially minimized and equilibrated for 1 ns; then, for 250 ps, the box was increased gradually along the *x* and *y* dimensions using different scaling factors. Therefore, at the end of this protocol, systems with different tension values were obtained, and the subsequent simulations at these surface tensions were performed keeping the *x* and *y* dimensions fixed.

The tubular bilayer (Figure 3C) was obtained from a DOPC vesicle built using Packmol (*51*). To this end, a central circular crown of 5 nm length was isolated and replicated 10 times. The replicas were then placed 1 nm apart and a subsequent unrestrained simulation led to the tubular structure by spontaneous self-assembly. This tubule was then equilibrated by using small holes to allow the adequate balance between the number of lipids in the inner/outer monolayers. The holes were created by applying a cylindric indenter of radius 3 nm acting on the beads that compose the PL hydrophobic talis. Then, the monolayers of a small portion of the tubule were separated in order to allow the insertion of a lens of 1794 TG molecules. The system was further equilibrated until complete reconstruction of the tubular structure, and the simulation was then run for additional 9.5 μs. We kept the holes open to maintain the lipids balance between the two monolayers.

To study the effect of inserting new TG molecules in the bilayer (Figure 2C), we slightly separated both PL monolayers along the *z* direction to accommodate more TG, and then those TG molecules outside the previously formed lens (see section *MD simulations analysis*) were replicated 3 times using the TopoTools VMD plugin(*52*).

To study the effect of unsaturation in TG chains on the equilibrium concentration of TG, systems containing 6050 DOPC and 1836 TOOP (a TG molecule with 2 oleic acids and 1 palmitic acid) or 1836 TOPP (a TG molecule with 1 oleic acid and 2 palmitic acids) were built.

### MD simulations analysis

TG lens formation was considered when an aggregate of at least 25 TG molecules within a 3.5 nm distance cutoff of a TG molecule was stable for at least 5 ns. The simulations were run until the formation of a TG lens was observed or for 1.5 μs if no aggregation occurred. For each TG concentration, bilayer composition, and setup, three independent simulations were performed. The rate of formation was calculated, from the average over the three replicas, as the inverse of the time of formation. Error bars were computed using error propagation as the standard deviation of the time of formation from the three independent simulations. For simulations where no formation was observed, the error bars were given as the inverse of the total simulation time (1.5 μs).

The analysis of the concentration of diluted TG was performed using the protocol outlined below. We defined a lens as a set composed by all the TG molecules found within a 5 nm distance of another TG molecule and not within 2.8 nm of a PL. The “lens-free bilayer” was accordingly defined as all lipid molecules that were at least 2.5 nm away from the lens. We tested different selections and we chose the selection at which the diluted TG concentration was converged (Figure S4) and that allowed us to consider the widest area of lens-free bilayer, excluding the side of the lens where TG molecules moved continuously inside and outside the lens (Figure S4). Except for the tubular structure, the simulations were run for 3 μs (see Table 1). The percentage of TG molecules inside the bilayer was obtained averaging the values of the last 1.5 μs of simulation, and the corresponding error bars were obtained from the standard deviation over two independent simulations. The dimensions of TG lenses were calculated using the minimum and the maximum *x*, *y* and *z* coordinates of the TG molecules at each side of the lens, averaged over time.

Trajectories of single TG molecules were obtained from a 500 ns trajectory of a system containing 11250 DOPC and 5508 TG using MDAnalysis(*53*) and plotted using MATLAB (Figure 1C-D).

Mean squared displacements (MSD) of TG and DOPC molecules were obtained using the LAMMPS command “compute msd” during production. Then, diffusion coefficients were calculated using the script msd2diff.py available in the ELBA-LAMMPS toolkit (https://github.com/orsim/elba-lammps/tree/master/tools).

The MSDs plotted in Figure S1 were obtained using the “Diffusion coefficient tool for VMD”(*54*).

For all simulations, surface tension was computed from the diagonal values of the pressure tensor (P_xx_, P_yy_, and P_zz_) using the Kirkwood-Irving method:

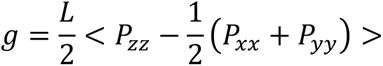

where L is the box length in the z dimension and <> means ensemble average.

### Yeast strains and growth conditions

Triple-mutant strain *pah1*Δ *are1*Δ *are2*Δ was generated by gene disruption, using PCR deletion cassettes and a marker rescue strategy(*55*). Quadruple-mutant strain, are*1Δ are2Δ dga1Δ lro1Δ,* was generated by mating and sporulation. The list of strains is shown in Supplementary Table 2.

Strains were cultivated in YP-rich medium (1% Bacto yeast extract, 2% Bacto peptone (USBiological Swampscott, MA)) or selective medium (0.67% yeast nitrogen base without amino acids (USBiological), 0.73 g/l amino acids) containing 2% glucose. Fatty acid-supplemented media either contained 0.24% Tween 40 (Sigma-Aldrich, St Louis, MO) and 0.12% oleic acid (Carl Roth, Karlsruhe, Germany) or 1% brij (Sigma-Aldrich, St Louis, MO) and 1mM palmitic acid (Sigma-Aldrich, St Louis, MO).

### Plasmids used in this study

To construct pTETO7-mScarlet, the DNA encoding m-Scarlet was synthesized (GenScript, Piscataway, NJ), PCR amplified and cloned into pCM189 plasmid(*56*) under TETO7 promoter. The plasmid expressing Pet10-mScarlet was constructed by amplifying *PET10* by PCR from *S. cerevisiae* genomic DNA and inserted into the plasmid pTETO7-mScarlet. The plasmid encoding the ER-DAG sensor was a gift from William Prinz(*29*) (National Institute of Diabetes and Digestive and Kidney Diseases, NIH, Bethesda).

### Fluorescence microscopy

Localization of Bodipy stained LD and mScarlet-tagged protein was performed by fluorescent microscopy of live yeast cells using a Visitron Visiscope CSU-W1 (Visitron Systems, Puchheim Germany). The Visitron spinning disk CSU-W1 consisted of a Nikon Ti-E inverted microscope, equipped with a CSU-W1 spinning disk head with a 50-µm pinhole disk (Yokogawa, Tokyo, Japan), an Evolve 512 (Photometrics) EM-CCD camera, and a PLAN APO 100× NA 1.3 oil objective (Nikon). To visualize LD in yeast, cells were grown in SC media and stained with Bodipy 493/503 (1 μg/ml) for 5 min at room temperature. Colocalization was evaluated by manually counting 100 lipid droplets stained either with Bodipy or the mScarlet-tagged marker protein. For *ole1^ts^,* we randomly chose 20 cells and manually counted LDs stained with Bodipy.

### Image rendering

All molecular dynamics images were rendered using VMD software(*57*) and graphs were generated using the software MATLAB.

Experimental images were treated using Image-J software and then resized in Photoshop (Adobe Photoshop CC 2015). Microscopic experiments were performed three times with similar results.

## Acknowledgments

We thank Vikram Reddy Ardham and Vineet Choudhary for useful discussions. This work was supported by the Swiss National Science Foundation (grant #163966). This work was supported by grants from the Swiss National Supercomputing Centre (CSCS) under project ID s726 and s842. We acknowledge PRACE for awarding us access to Piz Daint, ETH Zurich/CSCS, Switzerland. RS is supported by the Swiss National Science Foundation (31003A_17303) and the Novartis Foundation for medical-biological Research (19B140).

