## Supplementary Information for "Lipid droplet biogenesis is driven by liquid-liquid phase separation"

Figure S1. 2D mean squared displacement (MSD) of TG and DOPC molecules in flat lipid bilayers.

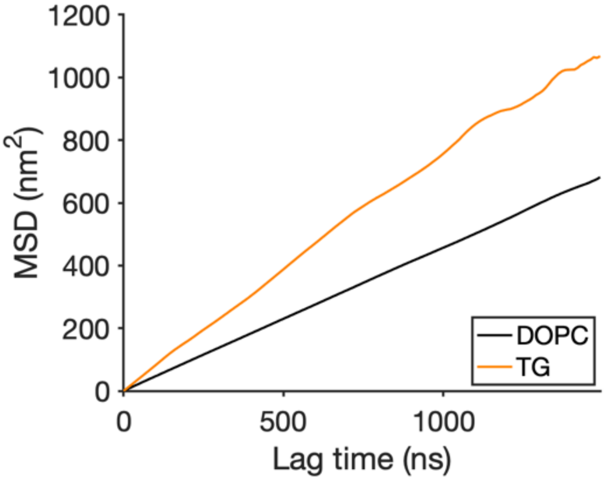


Table S1. Ratio between 2D diffusion coefficients ($D_{coeff}$) of TG and DOPC in different systems: (1) pure DOPC bilayer, (2) pure DOPC bilayer and tension=2 mN/m, (3) ER-like bilayer. The concentration of TG in all the systems is 2 % mol.

| System | Ratio $D_{coeff}$ TG/DOPC |
|---|---|
| (1) Pure DOPC | 1.6 |
| (2) Tension= 2 mN/m | 1.7 |
| (3) ER like | 2.0 |

Movie S1. TG molecules move freely in the bilayer and in the oil blister. Random molecules from the blister (red) and the bilayer (blue) are highlighted.

Movie S2. Oswald ripening effect. TG molecules from a blister of small size (blue) move toward bigger lenses due to difference of pressure in the three lenses.

Movie S3. TG lens dissolution in a DOPC bilayer. The concentration of TG is below (<1 mol%) the calculated "free TG" threshold in a DOPC bilayer(1.1±0.1 mol%).

Table S2. S. Cerevisiae strains used in this study

| Strain | Genotype/Description |
|---|---|
| RSY 3091 | *Matα his3Δ1 leu2Δ0 lys2Δ0 ura3Δ0 met15Δ0 ole1ts* |
| RSY 3077 | *Matα his3Δ1 leu2Δ0 lys2Δ0 ura3Δ0 met15Δ0 are1::KanMX are2::kanMX trp1::URA lro1::TRP dga1::Lox-HIS-Lox* |
| RSY 5165 | *Matα his3Δ1 leu2Δ0 lys2Δ0 ura3Δ0 met15Δ0 pah1:: KanMX are1::HIS3 are2::LEU2* |

Figure S2: Incorporation of saturated fatty acids in TG chains increases the accumulation of diluted TG in lipid bilayers.

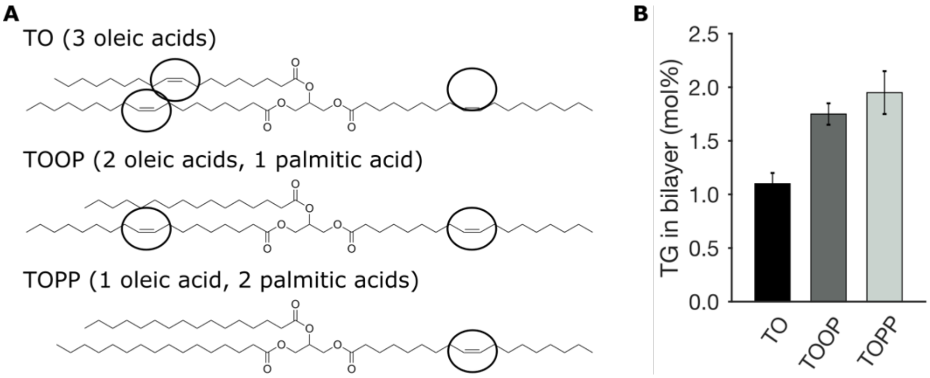


Figure S3: Comparison between the area per lipid in bilayers devoid of TG and the equilibrium concentration of TG in systems with the same bilayer composition.

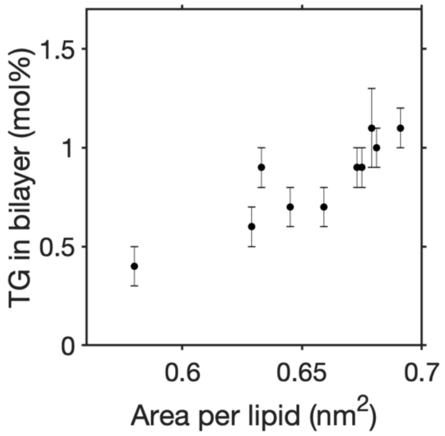


Figure S4. Choice of the radius for the calculation of diluted TG. We chose the smallest radius (black) above which we got always the same value of diluted TG.

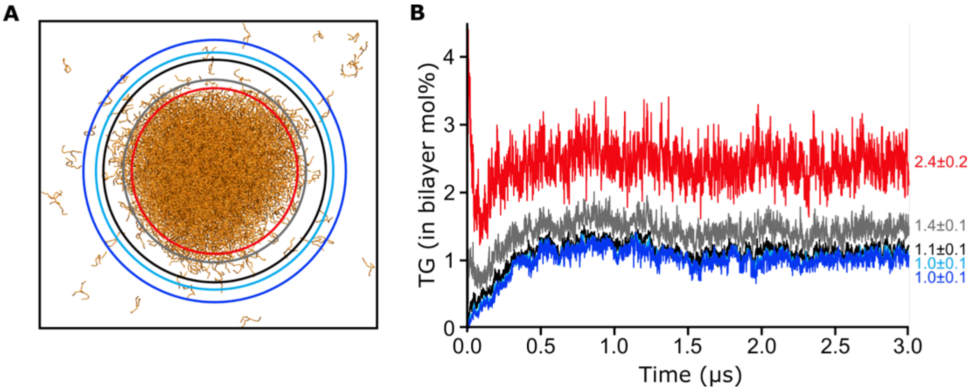
